# A thermodynamic scaling law for electrically perturbed lipid membranes: validation with the steepest-entropy-ascent framework

**DOI:** 10.1101/2020.12.03.410431

**Authors:** I. Goswami, R. Bielitz, S.S. Verbridge, M.R. von Spakovsky

**Affiliations:** Department of Bioengineering and California Institute for Quantitative Biosciences (QB3), University of California, Berkeley CA 94720; Department of Material Science and Engineering, University of California, Berkeley CA 94720; Department of Biomedical Engineering and Mechanics, Virginia Tech, Blacksburg VA 24061; Department of Mechanical Engineering, Virginia Tech, Blacksburg VA 24061

**Keywords:** Lipid membrane, Ising model, Steepest-entropy-ascent quantum thermodynamics, Thermodynamics, Pulsed electric fields

## Abstract

Experimental evidence has demonstrated the potential of transient pulses of electric fields to alter mammalian cell phenotypes. Strategies with these pulsed electric fields (PEFs) have been developed for clinical applications in cancer therapeutics, in-vivo decellularization, and tissue regeneration. Successful implementation of these strategies involves understanding how PEFs impact the cellular structures and, hence, cell behavior. The caveat, however, is that the PEF parameter space comprised of different pulse widths, amplitudes, and the number of pulses is very large, and design of experiments to explore all possible combinations of PEF parameters is prohibitive from a cost and time standpoint. In this study, a scaling law based on the Ising model is introduced to understand the impact of PEFs on the outer cell lipid membrane so that an understanding developed in one PEF pulse regime may be extended to another. Experimental study is used to argue for the scaling model. Next, the validity of this scaling model to predict the behavior of both thermally quenched and electrically perturbed lipid membranes is demonstrated via computational predictions made by the steepest-entropy-ascent quantum thermodynamic (SEAQT) framework. Based on the simulation results, a form of scaled PEF parameters is thus proposed for lipid membrane.

## 1. Introduction

Newer cancer therapeutic and tissue regeneration approaches have explored the idea of manipulating cell behavior *in-vivo* via the use of pulsed electric fields (PEFs) introduced into the tissue using a pair of electrodes. A representative clinically used PEF pulse train for soft tumor treatment (referred to in this article as μsPEF) consists of 80 to 100 square wave pulses each of width 100 μs delivered into the tissue at a frequency of 1 Hz. The μsPEF causes the formation of outer cell lipid membrane pores (i.e., a phenomenon termed electroporation) that either reseal or grow spontaneously contingent upon the pulse amplitude. ^1^Reversible electroporation in which pores ultimately reseal in time is clinically used for enhancing drug transport into the cytoplasm when the membrane is temporarily at a state of higher permeability. Irreversible electroporation (IRE) in which pores grow spontaneously leads to loss of cell homeostasis and ultimately cell death. IRE is clinically used in tumor tissue ablation and recently has emerged as a strategy for *in-vivo* tissue decellularization and tissue engineering [1]. Furthermore, the extent of the effect of μsPEF on the pro-cancerous inflammatory signaling is shown to be conditioned by pulse amplitude [2]. Other forms of PEFs incorporating lower pulse widths (≤ 1*μs*) but higher pulse amplitudes compared to μsPEF have been proposed for cancer therapeutic strategies as well. The permittivity of lipid membranes decreases, and conductivity increases as the frequency of an exogenous time-oscillating electric field is increased. Thus, exposing cells to lower pulse widths can allow the leveraging of differences in membrane properties at higher harmonics and targeting organelles. These sub-microsecond PEF modalities (i.e., nsPEF: nano-second PEF; H-FIRE: High Frequency IRE) have been used to elicit specific cellular responses such as, for example, targetting cancerous cells over their healthy counterparts [3].

The fact that different cellular responses can be achieved by altering PEF parameters is significant from a translational medicine point of view. However, this also raises a number of important mechanistic questions as to how PEFs influence cellular structures. This challenge is compounded by the vastness of the PEF pulse parameter space (i.e., the range of timescales for pulse width, the range of amplitudes, and the varying pulse numbers). Experimental and computational investigation of mechanistic questions for all combinations of PEF parameters (amplitude, pulse, and number) is not only cost prohibitive but likely impossible to carry out in a finite period of time. Attempts have been made to find equivalent pulses, i.e., develop mathematical relationships between pulse parameters, so that two equivalent pulses have similar outcomes such as the same fraction of electroporated cell population (e.g., [4, 5, 6]). The most extensive of these works in terms of the range of pulse widths covered (ns - ms) was reported by Puchihar *et al*. [4] whereas others were based on a very limited rangeof PEF parameter space. A few observations from these works is that the effect of pulses is not linearly additive (e.g., the limitations of linearity and the complexity of extrapolation have recently been reported by [7]), and it is difficult to derive a phenomological mathematical expression to cover the entirety of PEF pulse parameter space.

A more fundamental route to developing a scaling law,whereby the cellular compartment such as the cell membrane is represented by a Hamiltonian that captures some part of the system’s behavior and the effect of the PEF accomodated within the Hamiltonian promises to significantly reduce the complexity of representing the PEF parameter space. In this article, we propose a thermodyanamic scaling based run-on the Ising model for investigating PEF perturbed lipid membranes. Model lipid membranes or giant unilamellar vesicles (GUVs) have been used to experimentally study the effect of PEFs on the cell membrane [8]. These synthetic biomimetic systems allow investigators to study the direct effects of PEFs without the complexity of dileneating crosstalk from downstream processes that may be induced in a living cellular system. However, initial studies on GUVs have revealed electropores (i.e., pores in the membrane induced by the electric field) that are on the order of *μm* whereas electropores detected in cells exposed to PEFs are on the order of nm [9]. Thus, PEF parameter scaling models based on experiments performed on GUVs may not apply to cells unless a more fundamental route is taken. Our proposed scaling law is motivated by recent findings that the lateral reorganization of the lipid bilayers that lead to the formation of the “domains” involved in cell signaling can be described by the same Ising universality classes and scaling laws that apply to ferromagnetic materials near critical points ([10, 11, 12, 13]). In order to predict the non-equilibrium evolution of the lipid system, as would be the case in a transient perturbation due to PEF exposure, a novel computational approach based on the first-principle, non-equilibrium thermodynamic-ensemble framework called steepest-entropy-ascent quantum thermodynamics (SEAQT [14, 15, 16, 17, 18, 19, 20, 21, 22, 23, 24]) is used. We therefore demonstrate the suitability of the scaling law for PEF parameter space when studying lipid membranes and provide a discussion on how this approach may be extended to other cellular compartments.

## 2. Theory: Proposed scaling law and SEAQT

### 2.1. Ising model to represent electrically perturbed lipid membranes

Experiments, such as those by Blicher *et al*. [25], on lipid unilamellar vesicles show that the permeability of vesicles is highest close to the critical miscibility temperature (i.e. temperature at which the components are miscible and form a homogenous mixture). Blicher *et al*. show that the rate of pore formation in these lipid unilamellar vesicles is proporational to the excess heat capacity. Furthermore, Kraske and Mountcastle demonstrate that increasing the cholesterol content in vesicle membranes reduces the permeability of the vesicles to biomolecules [26], perhaps due to a change in the miscibility temperature [27]. The permeation rate of the lipid membranes shows a behavior similar to the heat capacity of an Ising model [25]. The Hamiltonian (i.e., an expression for the total energy of the system) for an Ising class, which describes phase transitions in ferromagnetic systems with magnetic dipole moments of atomic spins with +1 or −1 states, is given by

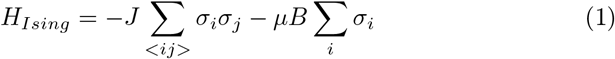

The first term *J* Σ_<*ij*>_ *σ_i_σ_j_* describes the interaction between nearest-neighbor spins, while the second term *μB* Σ_*i*_ *σ_i_* describes the interaction each spin has with an external magnetic field. In this equation, *J* is the interaction strength between neighboring spins *σ_i_* and *σ_j_* and is assumed to be constant between all pairs. The term *μB* represents the product of the magnetic moment *μ* and the applied external magnetic field *B*. According to the conventional Ising model, the summation of the spins (Σ_*i*_ *σ_i_*) is termed magnetization. The absolute value of the ensemble-averaged magnetization being close to 1 implies homogeneity of phase or in other words the dominance of one of the two spins over the other. The ensemble-averaged magnetization being close to 0 implies a heterogenous state in which there is equal probability for either spin. Interestingly, such Ising-type behavior has also been reported in lipid membranes without any interaction with an external field [11, 13].

We questioned whether PEF induced poration can be viewed as a a phase change problem whereby the poration is a maximum at a critical point dictated by the miscibility point (note that this miscibility point is a function of both the local composition as well as the temperature). We then investigated whether cholesterol content in the cell membrane can be a gross parameter that can dictate susceptibility to PEF induced poration. Comparison between the membrane cholesterol contents of non-cancerous 10A, ductal carcinoma insitu DCIS.com, and triple negative breast cancer MDA-MB-231 cell lines show that the more aggresive MDA-MB-231 has approximately 25% higher values of cholesterol (**Figure 1A**). Lower cholesterol content corresponded to higher susceptibility to a μsPEF as measured by the viability of the cell population after exposure (**Figure 1A**). Furthermore, as has been shown previously for nsPEF [28], we see an increased susceptibility of the MDA-MB-231 cells to a PEF when the cholesterol content of the membrane is pharmacologically reduced with acute exposure to methyl-β-cyclodextrin (MβCD; see **Figure 1B**). These observations are analogous to the ones made by Kraske and Mountcastle, albeit with the caution of not over generalizing the importance of cholesterol in PEF susceptibility.

**Figure 1:**
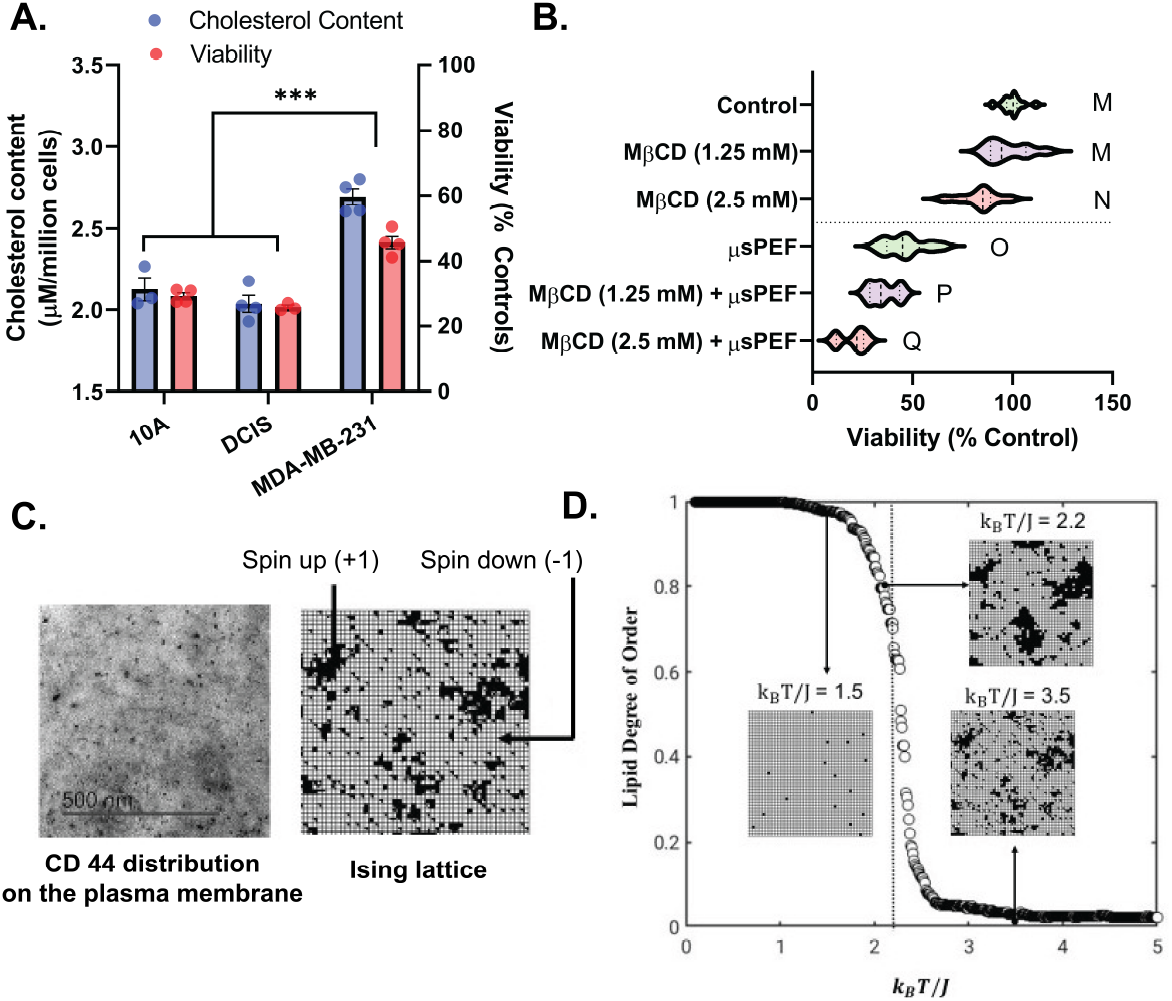
Lower cholesterol content in the cell membrane in cell lines 10A and DCIS compared to MDA-MB-231 corresponds to higher susceptibility to the μsPEF **(A)**. Reduction of membrane cholesterol in the MDA-MB-231 cell line via the MβCD treatment increases the susceptibility to the μsPEF **(B)**. Statistically non-distinct conditions are grouped by letter (e.g., M vs. M), whereas statistically distinct conditions are grouped by different letters (e.g., M vs. O) to illustrate significant differences as determined by a one-way ANOVA followed by a Tukey’s HSD posttest. The lipid domains on membranes are highly ordered regions. A visualization of the distribution of CD44 protein receptors localized in these ordered regions is shown in an electron microscopy image **(C)**. The lipid cell membrane is represented using an Ising lattice of spins up (black) and spins down (white) **(C)**. Equilibrium predictions of the absolute value of the normalized lipid degree of order is obtained via the Metropolis scheme on an Ising Hamiltonian and describes phase behavior on either side of the scaled critical parameter, in this case temperature, as shown in **(D)**.

The arguments presented above and the observation that the lipid membrane has an Ising-type behavior forms the basis of the scaling law proposed here for PEF parameters and is used to study the impact that electric fields have on a lipid membrane. Accordingly, the proposed Hamiltonian for the Ising model is given as

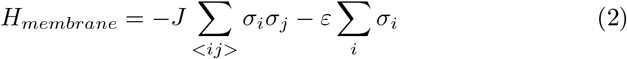

In this model, the term *ε* is the scaled PEF term for a given lipid membrane system. Normalization with the interaction term reduces **Eq. 2** to *H_membrane_/J* = −∑_<*ij*>_ *σ_i_ σ_j_* – (*ϵ* / *J*)∑_*i*_ *σ_i_*. The term *ϵ/J* is the universal equivalent scaled PEF parameter. While the form of this scaled PEF term will be discussed later, this term does not contain the number of pulses. The non-linearity arising from multiple pulses is taken into account by the behavior of the Ising model itself as is seen later in the manuscript.

The spins of −1 and +1 in this model represent lipid liquid-ordered and liquid-disordered phases, respectively. Note that the selection of which spin represents which phase is not important. Ordered phases are associated with sub-micron sized structures or “domains” that are enriched with certain lipid and protein species. Such cholesterol-rich domains can be visualized using electron microscopy. In **Figure 1C**, electron microscopy is used to visualize the distribution of the protein CD44 that has been demonstrated to exist in large numbers in lipid domains [29, 30, 31, 32, 33]. Thus, a membrane may be represented mathematically as an Ising lattice of spins down (−1) and spins up (+1) as shown in **Figure 1C**. Since the Ising model proposed here is used to capture the effects of PEFs on domain formation in lipid systems, the magnetization term (Σ_*i*_ *σ_i_*) becomes the lipid degree of order and represents the dominance of one or the other of the spins or phases. Below a critical point, the lipid membrane is dominated by a single phase or spin, and, therefore, the absolute value of the lipid degree of order is close to 1. In **Figure 1D** (the plot of the absolute value of the normalized lipid degree of order versus (*k_B_T/J*)), this critical point occurs at approximately *k_B_T/J* = 2.3 when the temperature alone is the critical parameter. Above the critical point, the lipid membrane is highly heterogeneous and, therefore, has a lipid degree of order close to zero.

To validate whether or not this scaling model is suitable for capturing the effects of PEFs on lipid membrane domain formation, computational predictions of the transient behavior of a perturbed lipid membrane are needed so that they can be compared with the experimental data developed in this study. The SEAQT framework is used here to do this. The rationale for using this framework is provided in the next section.

### 2.2. SEAQT framework and equation of motion for lipid membrane

To validate the scaling model, the non-equilibrium, ensemble-based SEAQT framework is used. It is able to model systems undergoing non-equilibrium processes, even those far from equilibrium, from the atomistic to the macroscopic level[23, 34, 24, 35, 36, 37, 38, 39, 40, 41, 42, 43, 44, 45, 46, 47, 48, 49]. This framework is not limited to any type of state (i.e. nonequilibrium, steady, unsteady, or metastable, unstable or stable equilibrium) or system size (i.e., a single atom or bulk sample) since it does not utilize intensive properties (e.g., temparature, pressure, chemical potential, etc.) as its state variables. Furthermore, the entropy employed is the von Neumann entropy, which has its basis a quantum or “quantum inspired” (e.g. Ising, Potts, or Heisenberg model) description. Thus, both quantum and classical systems can be treated within this framework. In addtion, SEAQT uses a density or so-called “state” operator 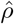 represents the state of the system, and the evolution of this density operator is tracked in time using the following SEAQT equation of motion (EOM):

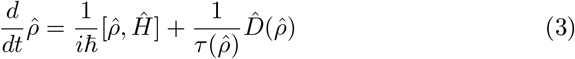

where the first term to the right of the equal sign captures the reversible symplectic dynamics (sub-manifolds on which purely Hamiltonian evolution takes place [50]), while the second term captures the non-linear dynamics of the irreversible relaxation of state. This second term takes the form of a dissipation operator 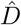 [18, 19] determined by a constrained gradient in Hilbert space and constructed on the basis of the principle of steepest entropy ascent [21] using a set of operators each of which is associated with one of the conservation laws to which the system is subjected. Note that the equation with only the first term on the right-hand side is the time-dependent Schrodinger (or equivalent von Neumann) equation, while the second term describes the non-linear dynamics characterized by entropy generation. In **Eq. 3**, 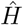 represents the Hamiltonian operator and *τ* is a relaxation time (a functional of the state operator or a constant) that can be determined from experimental data (e.g., [41, 34, 47, 43]) or in a completely *ab initio* manner (e.g., [21, 24, 36, 37]). The kinetic behavior of the system (i.e., the unique thermodynamic path) predicted by the EOM is independent of *τ*, which simply captures the dynamics of state evolution, i.e., the time required to traverse the thermodynamic path. Thus, *τ* links the dynamics to the kinetics via a corrected timescale. The density operator 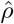 is used to obtain the expectation value (represented by 〈〉) of any extensive property *P* of the system given an operator 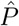, i.e.,

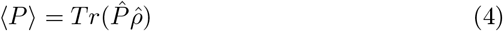

The SEAQT framework is used below to provide unique insights into lipid systems perturbed thermally and electrically.

The general EOM as described by **Eq. 3** is formulated specifically for a lipid membrane represented by an Ising lattice. Determining the eigenstructure of this lattice, whose energy is represented by the Hamiltonian *H_membrane_* of **Eq. 2**, involves solving an energy eigenvalue problem of the form

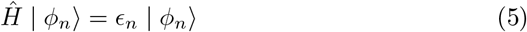

where 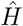 is the corresponding operator form of *H_membrane_*. The former is not known explicitly and, thus, **Eq. 5** cannot be solved directly. Instead a stochastic approach as described in section **3.4** and **Appendix A.2** is used to determine the |*ϕ_n_*), which are the eigenvectors of the eigenstructure. The *ϵ_n_* are the eigenenergies, and each paired |*ϕ_n_*) and *ϵ_n_* represents a possible eigenstate of the system. Following the terminology used in linear algebra, *ϵ_n_* is named an eigenlevel (or eigenvalue) and the set of energy eigenlevels that encompasses all possible eigenstates of the system can be used along with their respective eigenvectors to construct the Hamiltonian operator 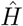 such that

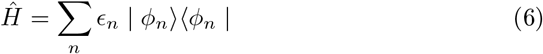

The thermodynamic state of the system at any instant of time is then some combination of available eigenstates determined by a distribution of probabilities or statistical weights among these levels. Note that the eigenlevel probability does not depend on whether the system is at stable equilibrium or not at stable equilibrium. In the case of the former, the probability for each eigenlevel is given by a canonical ensemble (i.e., one for which the volume and the number of particles constituting the system are fixed) expressed as

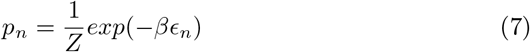

where *β* is the inverse of the product between the Boltzmann constant (*k_B_*) and the thermodynamic temperature (*T*) at stable equilibrium (i.e., *β* = 1/*k_B_T*). The partition function *Z* = ∑_*i*_ *exp*(−*βϵ_i_*) provides the normalization condition, i.e., 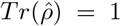. The diagonal terms of the density operator 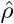 contain the probabilities associated with each energy eigenlevel at a given state provided the density operator is diagonal in the energy eigenvalue basis of the Hamiltonian.

As indicated in Li and von Spakovsky [24], for many classical systems in which there are no quantum correlations and the choice of Hilbert space metric used to construct the dissipation operator 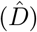 is the Fisher-Rao metric, the density operator is indeed diagonal in the energy eigenvalue basis so that 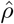 and 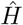 commute and the first term to the right of the equal sign of the EOM (**Eq. 3**) vanishes. The EOM then reduces to an irreversible relaxation only that can be expressed as the probabilities associated with each eigenlevel evolving in time via

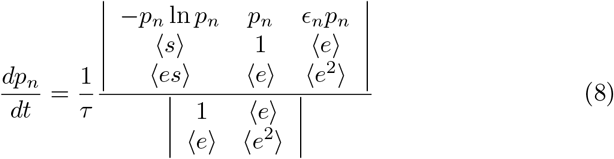

Here 〈·〉 represents the expectation value of a thermodynamic property, and the only generators of the motion are the identity operator 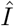 and 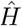 [18]. As mentioned earlier, the expression for the SEAQT EOM contains the thermodynamic entropy (〈*s*〉) and energy (〈*e*〉) which are defined for both equilibrium and non-equilibrium states (or more generally, not stable equilibrium states [51]).

In order to predict the evolution in state of the lipid membrane as represented by the Ising Hamiltonian, all the possible energy eigenlevels that the system may encounter must be determined. This task is not trivial. If one considers an Ising lattice of ‘N’ nodes with each node having an equal probability of being occupied by a spin up (+1) or spin down (−1), there are 2^*N*^ possible lattice configurations. Therefore, theoretically there can be 2^*N*^ energy eigenlevels. However, to complicate matters, it turns out that the energy eigenlevels are degenerate (i.e., multiple lattice configurations (eigenvectors) have the same energy), and, therefore, the number of energy eigenlevels is much less than 2^*N*^. The degenerate eigenlevels and eigenvectors can, therefore, be grouped together, and the EOM (in **Eq. 8**) can be re-stated as

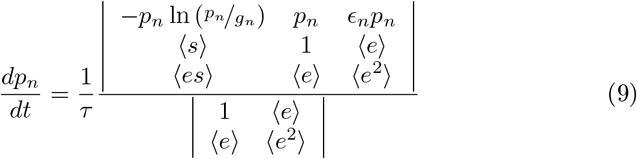

where the term *g_n_* denotes the degeneracy associated with the *n^th^* eigenlevel and *τ*\* is the dimensionless time defined by the ratio of the actual time and the relaxation time (*τ*\* = *t*/*τ*) provided *τ* is a constant and not a function of 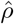. If it is not constant, then 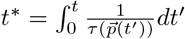. **Eq. 9** is used in this study to simulate the lipid dynamics of electrically perturbed lipid membranes. The next section describes both the experimental and numerical methods used to generate the experimental data and make the numerical predictions.

## 3. Methods

### 3.1. Cell line and culture conditions

The cell lines MCF-10A and ductal carcinoma in-situ MCF-DCIS.com were obtained from Dr. Eva Schmelz at Virginia Tech (Blacksburg, USA). The human triple negative breast cancer cell line MDA-MB-231 was purchased from ATCC (Catalog number: HTB-26). The MDA-MB-231 cells are maintained in a DMEM-F12 culture medium supplemented with 10% (by volume) fetal bovine serum and 1% (by volume) penicillin streptomycin. The MCF-10A and MCF-DCIS.com cell lines are grown in a DMEM-F12 culture medium supplemented with 5% (by volume) horse serum, 20 ng/mL endothelial growth factor, 0.5 mg/mL hydrocortisone, 10 mg/mL insulin, and 1% (by volume) penicillin streptomycin. Cells are sustained in humidified incubators at 37 °C and 5% CO_2_. Cells are sub-cultured at approximately 80% confluence, and a 0.25% Trypsin-EDTA solution is used for detachment. All experiments are performed within the first ten sub-cultures.

### 3.2. Pulsed electric field delivery

The cells are seeded into a 12-well tissue culture plate with ~50,000 cells per well. Cells are allowed to reach ~80-90% confluency. Before PEF exposure, the cells are washed gently 3x with PBS, taking care so as to not shear the cells from the bottom of the plate. 1 mL of RPMI-1640 basal medium (a medium without serum and penicillin streptomycin) is added to each well. A simplified illustration of the experimental setup is shown in **Figure Appendix A.1**. PEF treatment is delivered to the cells using a modified tissue culture lid. Briefly, a 1/16th drill bit is used to drill holes that are 2.0 cm apart along the center line of each column. Disposable stainless-steel biopsy needles, 1.6 mm in diameter, are used as electrodes and are introduced into the wells at diametrically opposite extremities of the well and placed in contact with the the bottom of the culture plate. Electrodes are connected to the BTX Harvard apparatus ECM630 electro cell manipulator generator via alligator clips. A clinically used pulse train with an applied voltage of 2000 V, a pulse widthof 100 μs, 99 pulses in the train, and a pulse interval of 1 sec is administered. Untreated controls are shams with 0 V. After all conditions have been pulsed, cells are returned to the humidified incubator at 37 °C with 5% CO_2_ for 3 hr and then washed 3x with 1xPBS. Cells are given a growth medium and returned once more to the humidified incubator and allowed to sit overnight (12 – 16 hours) before further processing and analyses.

### 3.3. Assays

#### 3.3.1. Viability assay via resazurin metabolic measurements

Prior to treatment, a solution of 10% by volume Alamar Blue solution is prepared using supplemented growth medium (refer to the cell lines and culture conditions). Baseline metabolic activity measurements are taken using a 10% Alamar Blue solution to assess cellular viability. 1mL of Alamar Blue solution is placed in each well. Sampling is done at four different intervals (*t* = 20 min, 40 min, 60 min, and 90 mins) with 100 μL drawn at each time point per well and transferred into a 96-well plate. A 96-well plate reader is used to measure baseline fluorescence values. For viability measurements, Alamar Blue readings are taken the following day, as initially described. Alamar Blue viability is assessed as a percentage of the fluorescent values after treatment in relation to the average baseline control measurements.

#### 3.3.2. Membrane cholesterol quantification

The cells are subcultured and seeded into a 6-well tissue culture plate at approximately a million cells per well using a hemocytometer to approximate the cell count. Cells are first washed with PBS and then acute extraction of membrane cholesterol is performed via a treatment with methyl-β-cyclodextrin (MβCD). Supernatant is then collected and analyzed for cholesterol using the Amplex Red Cholesterol Assay Kit (Invitrogen catalog A12216) following the standard protocol provided by the manufacturer.

### 3.4. Estimation of the density of states

The information about the energy eigenlevels and the degeneracy associated with each eigenlevel are obtained via two different methods: the Wang-Landau algorithm (WLA) [52, 53] and the Ren-Eubank-Nath (REN) algorithm [54].

Brief details of their implementation are provided in **Appendix A.2** and more detailed derivations are given in [55].

### 3.5. Solving the SEAQT EOM

The **Eq. 9** is used to capture the behavior of a lipid membrane represented by an Ising Hamiltonian. The system of non-linear ordinary differential equations formed by the EOM is solved using the ODE15s and ODE45 solvers available in the MATLAB (Mathworks, Natick MA, USA) programming software.

## 4. Results

### 4.1. Lipid membrane eigenstructure and the SEAQT predicted relaxation

To determine the eigenstructure of the lipid membrane, represented by the Ising Hamiltonian, the eigenvalue problem given by **Eq. 5** is solved via statistical techniques, i.e. the WLA and REN algorithms (refer to sections 3.4 and Appendix A.2). Both algorithms provide a way to extract system information in terms of the energy eigenlevels and their respective degeneracies. **Table 1** compares the number of distinct eigenlevels for the univariate (i.e., without the field) and bivariate (i.e., with the field) Ising Hamiltonians. While the number of distinct eigenlevels for a univariate case varies linearly with respect to the number of nodes 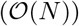, the variation for the bivariate is non-linear 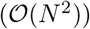. The eigenstructure of a system represented by a bivariate Ising Hamiltonian is shown in **Figure 2A**. This eigenstructure information allows one to determine the degeneracy (on a logarithmic scale) for each combination of the interaction term 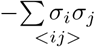 and the lipid degree of order 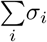 corresponding to a unique eigenlevel. Each discordant edge number corresponds to a constant value of the interaction term 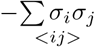. The sum of the degeneracies along a discordant edge number correspond to the degeneracy associated with an eigenlevel in the univariate distribution. Thus, the bivariate prediction collapses to the univariate predictions as shown in **Figure 2B**.

**Figure 2:**
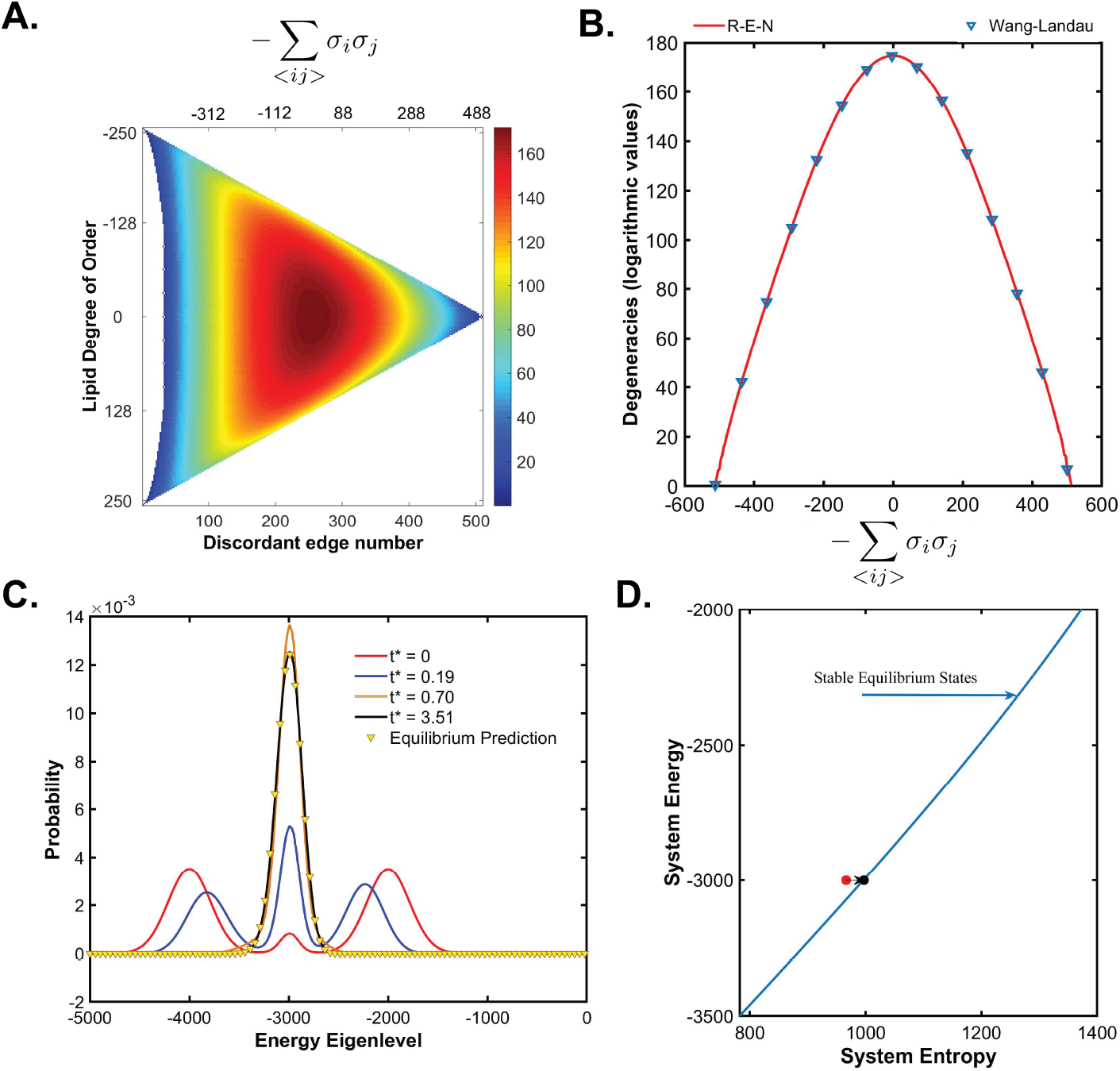
Eigenstructure of the lipid membrane, i.e., the information about all possible eigenlevels and their respective degeneracies are predicted using statistical techniques (**A-B**). Eigenlevels and their respective degeneracies (on a logarithmic scale) for a 16 x16 lattice represented by a bivariate hamiltonian (**A**). An eignestructure for a univariate Ising Hamiltonian for a 16×16 lattice (**B**). The SEAQT EOM uses the eigenstructure information to predict the evolution of the probabilities of the eigenlevels of the lipid membrane relaxing from a state of non-equilibrium (−) to equilibrium (−; **C**) shown here for a 50×50 lattice. Evolution of the lipid membrane system entropy as the system state relaxes to an equilibrium state is visualized on an energy vs. entropy map (**D**). Note that the units of energy and entropy are dimensionless here since they are normalized to *J* and *k_B_*, respectively.

**Table 1:** Number of distinct eigenlevels in an Ising lattice.

|  | *Univariate*<br>(Hamiltonian without field) | *Bivariate*<br>(Hamiltonian with field) |
| --- | --- | --- |
| $2 \times 2$ | 3 | 6 |
| $4 \times 4$ | 15 | 80 |
| $6 \times 6$ | 35 | 482 |
| $16 \times 16$ | 255 | 29,430 |
| $32 \times 32$ | 1,023 | 497,260 |
| $50 \times 50$ | 2,499 | 3,021,498 |
| $100 \times 100$ | 9,999 | $> 10^7^*$ |

*\*Extrapolated using a curve fit between the number of eigenlevels vs. the number of nodes ( $N=4$ through $N=2500$ )*

Once the eigenstructure is obtained, information regarding the degeneracies and eigenlevels required in **Eq. 9** is obtained. With this information it is possible to prepare a system in an arbitrary initial non-equilibrium or equilibrium state (see section Appendix A.3 for more details on the preparation of initial states and on details of the computation). A validation is carried out to investigate whether or not the SEAQT framework can predict the relaxation of an isolated system represented by the Ising Hamiltonian from a state of nonequilibrium to that of stable equilibrium. An isolated system has no boundary interactions, and, therefore, the expectation value of the system energy remains invariant. The relaxation process from non-equilibrium to equilibrium results in a spontaneous increase in system entropy due to entropy generation as would be expected based on the second law of thermodynamics. The relaxation of the isolated Ising system, predicted by the SEAQT EOM, is shown here both for the univariate distribution (i.e., tracking only the probabilities associated with the energy eigenlevels; see **Figure 2C-D**) as well as the bivariate distribution (i.e., tracking the probabilities associated with both the energy eigenlevels as well as the lipid degree of order; see **Figure A.2**). The solution of the SEAQT EOM for the state evolution of the Ising lattice provides the temporal evolution of the state of the thermodynamic system relaxing from a state of non-equilibrium to equilibrium, and its prediction of the equilibrium state compares well with the equilibrium predictions obtained from WLA. In addition to tracking the system’s state in the energy-entropy plane, the bivariate predictions allow the temporal tracking of the system’s lipid degree of order as seen in **Figure A.2A-C**. This is useful in modeling lipid membranes perturbed with PEFs.

### 4.2. Thermal quenching of the lipid membrane

While the previous section successfully demonstrates the ability of the SEAQT framework to predict the non-equilibrium and equilibrium states of an isolated Ising system representing a lipid membrane, it is important to demonstrate that an interaction a with PEF and thermal reservoir can also be modelled. For the latter, the focus here is on replicating the scenario of lipid membrane quenching performed by Veatch *et al*. [11] and Honerkamp-Smith *et al*. [13, 12] to arrive at the conclusion that lipid membranes have the same critical behavior as ferromagnets. This is important in deriving a scaled PEF parameter as will be discussed in section 5.

Veatch *et al*. [11] performed quenching experiments on giant plasma membrane vesicles (GPMVs) stained with DilC12 or fluorescent antibodies to track the protein receptor *Fcϵ*RI. The GPMVs are placed between two glass coverslips and observed under a fluorescent inverted microscope. Next, the temperature of the system consisting of the GPMVs is either increased or decreased in steps and the fluorescence of the lipid membrane measured after allowing 5-10 min of equilibration following a temperature step. Spatial organization of the lipids on the GPMVs are traced and images post-processed to obtain statistical measures such as correlation lengths and line tensions. They report that all GPMVs undergo a phase change whereby at temperatures above a miscibility temperature (recorded in the neighborhood of ~ 20°C) the GPMVs are heterogenous with ordered and disordered phases seemingly appearing as a single phase. Below the miscibility or critical temperature, the GPMVs experience phase separation. Note that the value of the critical temperature (*T_c_*) is associated with the critical point of the Ising system given by *k_B_T_c_*/ *J* = 2.3.

To replicate the results of the Veatch *et al*. experiment and to demonstrate the computational capability of the SEAQT framework to capture the Ising-like behavior of the bio-membrane, the membrane represented by an Ising lattice is modeled interacting with a thermal reservoir. Li and von Spakovsky introduce the hypo-equilibrium concept [23] and use it to generate a specific SEAQT EOM for a system undergoing an interaction with a thermal reservoir [24]. If a system interacts with a large thermal reservoir such that the specific heat of the reservoir is much larger than that of the system (i.e., *C_reservoir_* ≫ *C_system_*), then the EOM represented by **Eq. 9**) reduces to

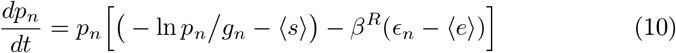

where *p_n_, g_n_*, and *ϵ_n_* represent the probability, degeneracy, and energy, respectively, associated with each eigenlevel of the system. 〈*s*〉 and 〈*e*〉 are the expectation values of the entropy and energy, while the term *β^R^* is inversely proportional to the temperature of the thermal reservoir (*β^R^* = 1/*k_B_T_R_*). The system is prepared in a state of stable equilibrium and then allowed to interact with the reservoir. The probabilities associated with the eigenlevels are then predicted using the SEAQT EOM, **Eq. 10**.

**Figure 3A-C** provides a representative prediction made by the SEAQT model of lipid membrane quenching for the univariate case. The membrane is represented by a 50×50 lattice. The system is initially at a stable equilibrium state where the value of *k_B_T*/*J* = 3.5 and is then allowed to interact with a thermal reservoir at *β^R^* = 0.5 (i.e., *k_B_T_R_* = 2). Interaction with the reservoir leads to a shift of the probability distribution towards energy eigenlevels associated with lower temperatures (**Figure 3A**). The membrane undergoes the process seen on the energy-entropy plot in **Figure 3B**. Note that although it follows a path of successive equilibrium states it is nonetheless an irreversible path and not a quasi-equilibrium (i.e., reversible) path since movement along the path is not restricted to very slow times. Interestingly, upon arrival at the critical point (*t*\* = 0.084 in **Figure 3A**), two peaks of the probability distribution appear, one on either side of the critical composition. At this critical point, the membrane is thrown out of equilibrium as seen in **Figure 3B** (albeit still close to equilibrium). The probability distribution of the energy eigenlevels converges to that of the stable equilibrium state at *k_B_T* / *J* = 2 at which point the system comes to its final state in the process. Since the process follows a sequence of equilibrium states, the system lipid degree of order as expected follows the same behavior as that of the equilibrium values predicted by the Metropolis-Hastings scheme (**Figure 3C**).

**Figure 3:**
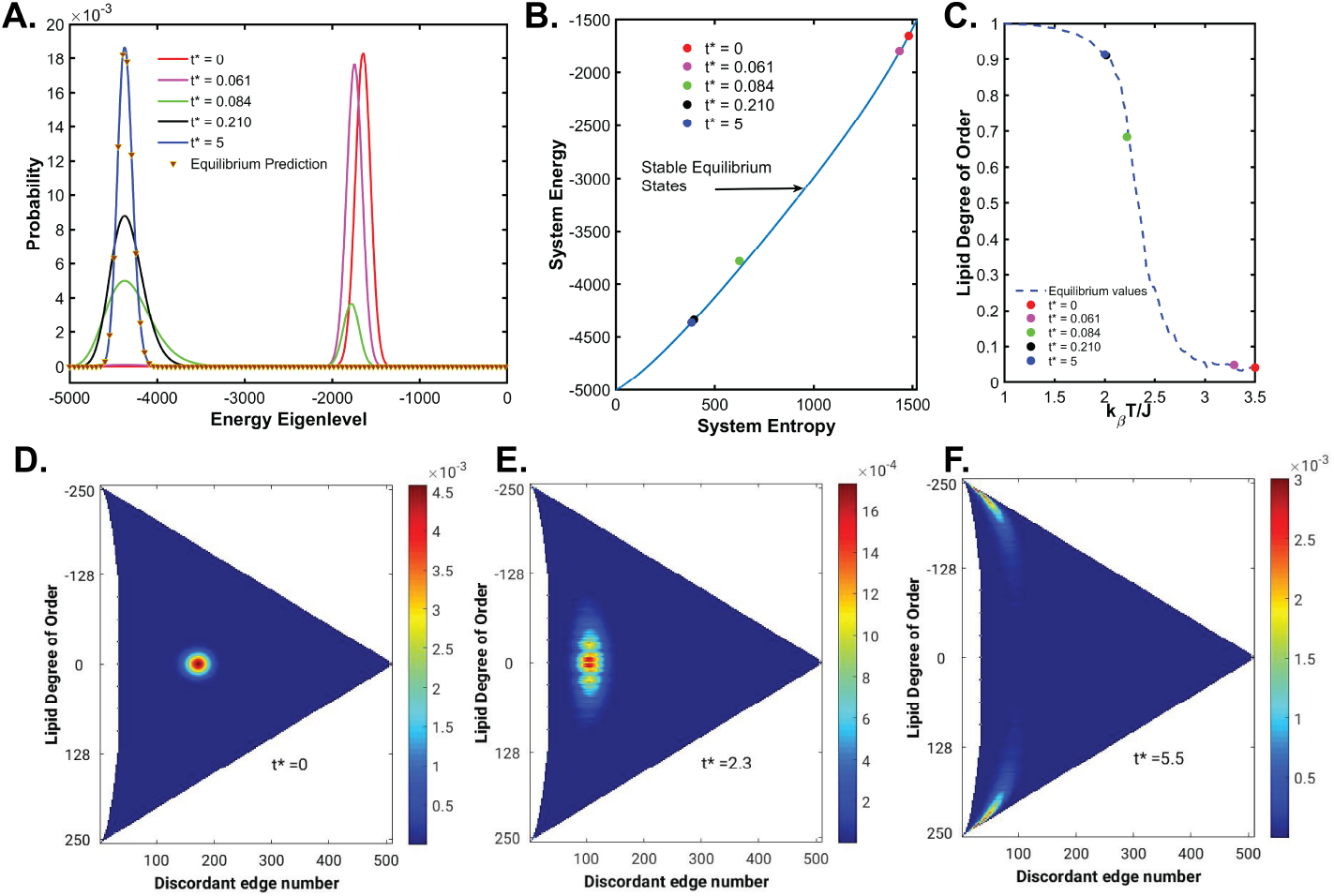
The evolution of the energy eigenlevel probabilities of a thermally quenched lipid membrane represented by a univariate Hamiltonian is shown in (**A**). The lipid membrane follows a sequence of equilibrium states as seen by the traces along the stable equilibrium curve (**B**). The expected lipid degree of order is shown in (**C**). The time-frames shown in (**D-F**) capture the evolution of the bivariate probability distribution as the lipid membrane is quenched. Starting at a state with an expectation value for the lipid degree of order = 0 (**D**), the system finally equilibrates with the reservoir temperature and eigenstates are concentrated towards the two extreme ends of the lipid order implying phase segregation or homogeneity (**F**).

To test whether or not the alteration in lipid degree of order is captured in the bivariate predictions of the SEAQT model, a simulation is performed to investigate the bivariate evolution of a 16×16 lattice interacting with a thermal reservoir. A smaller lattice size of 16×16 for the bivariate case is chosen here simply to minimize the computational costs for this illustration. For the bivariate case, an initial state is prepared such that the membrane is initially at a stable equilibrium state where *k_B_T*/ *J* = 3.5 with the expected value of the lipid degree of order equal to zero (**Figure 3D**). The membrane is then allowed to interact with a thermal reservoir at *β^R^* = 0.4543 (i.e., *k_B_T_R_* = 2.201) and the probability distribution evolution predicted using the SEAQT EOM **Eq. 10** is seen in **Figure 3D-F**. As in the univariate case, the bivariate predictions show the membrane evolving towards the energy states associated with lower temperatures. Until the system arrives at the critical point, the membrane maintains a lipid degree of order that equals zero. At and below the critical point, a splitting is seen in the probability distribution (e.g., **Figure 3F**) with a dichotomy between the eigenstates with opposite values of the lipid degree of order. Thus, below the critical point the probability distribution has two separate branches each located at one extreme end of the lipid degree of order. This represents a phase segregation in the membrane. Note that the choice of *β^R^* in both sets of simulation is based on the fact that they represent values below the critical point of the Ising model (i.e. *k_B_T_c_*/ *J* = 2.3).

The SEAQT EOM correctly predicts the phase transition associated with lipid membrane quenching. Note that in this section the phase transition from disordered to ordered states is brought about by a reduction in temperature below a critical temperature. However, a phase transition may also be brought about via a change in the composition. Thus, the critical point is a function of both the temperature and composition, and this point is to be kept in mind for the upcoming section, which is focused on understanding how PEF pulse parameters influence the order of the lipid membrane. As will be discussed in subsequent sections, this has implications for cell membrane criticality related to cell signaling.

### 4.3. Predicting lipid membrane behavior in the presence of a PEF

In this section, the SEAQT framework is used to simulate lipid dynamics perturbed with a PEF. We explore the meaning of the scaled PEF parameter term *ε* as well as the dimensionless time of exposure *τ_pw_* and discuss how such a scaling term may be derived using experiments. Choosing the initial state for the simulation is a challenging task since one has to justify its relevance to experiments. It is suggested here that the initial condition be prepared to represent two scenarios. First, an ordered state found below the critical miscibility point **Figure A.3A** is considered. Next, a state with a non-zero lipid degree of order that is found above the critical miscibility point is used. As shown in **Figure A.3B**, such a state of non-zero lipid order is not favored at equilibrium. It is important to realize that the critical point is a function of both composition and temperature. While the criticality in the plasma membrane of living systems such as mammalian cells is achieved via alteration in composition (since temperature is constant), from the perspective of an experimental setup with GPMVs/GUVs, a thermodynamically favorable initial state is the only viable condition. As will be discussed later, experimental findings from the GPMVs/GUVs can be extrapolated to living systems as has been shown by [11, 13]. Thus, the discussion on the effect of PEF parameters is focuses on this thermodynamically favored intial state, i.e., a preparation of the lipid membrane below the critical point at a constant temperature. Note that all the results in this section are obtained from the bivariate predictions of a 16×16 lattice.

#### 4.3.1. Effect of the scaled parameter term on lipid membrane criticality

A typical PEF pulse train used in this section is shown in **Figure 4.3A**. First, the lipid membrane is exposed to a PEF pulse with constant *ε* = 1 but for varying times (*τ_pw_* = 0.125,0.25,0.5), i.e.,

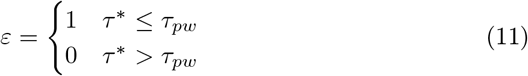

**Figure 4:**
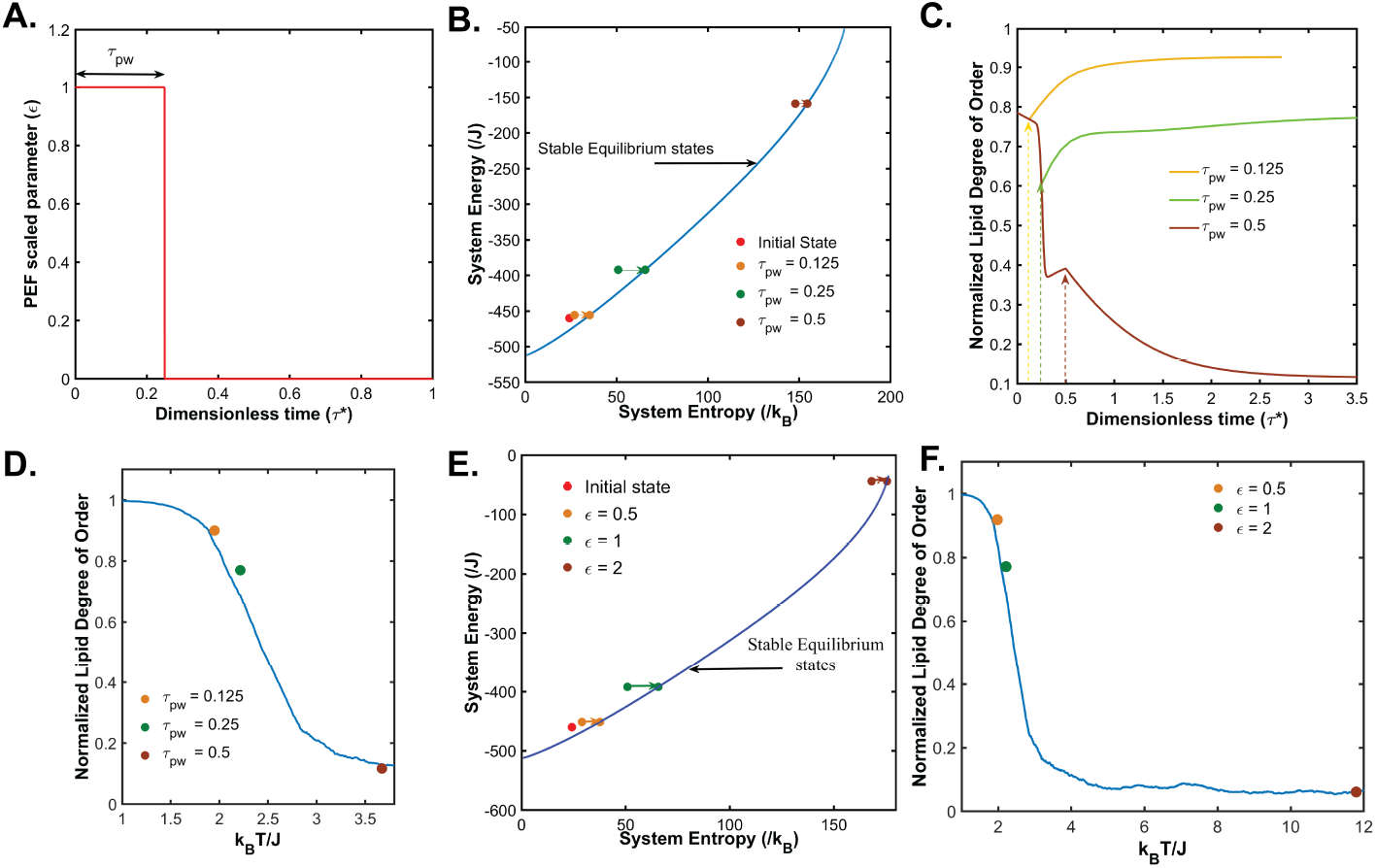
Effect of the PEF on the lipid membrane. The lipid membrane is exposed to the PEF pulse represented in (**A**). SEAQT predictions of the state of the lipid membrane treated with varying exposure times *τ_pw_* (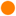: 0.125; 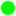: 0.25; 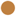: 0.5; **B-D**) and the scaled PEF parameter *ϵ* (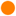: 0.5; 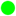: 1; 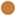: 2; **E-F**) of a single PEF pulse. The system is tracked in the energy-entropy plane (*τ_pw_*: **B**; *ϵ*: **E**). In these energy-entropy representations, the state of the membrane immediately after the pulse is in non-equilibrium but is allowed to relax (as shown by the direction of the arrows) towards equilibrium. Normalized lipid degree of order of the membrane as predicted by the SEAQT EOM during the PEF exposure is shown for *τ_pw_* in (**C**). The states corresponding to equilibrium after PEF exposure predicted by the SEAQT EOM are plotted relative to the lipid order versus the *k_B_T*/ *J* curve for the equilibrium values predicted by the Metropolis-Hastings scheme (*τ_pw_*: **D** and *ϵ*: **F**).

The evolution of probabilities in dimensionless time is predicted using the SEAQT EOM (**Eq. 9**). The final probability distribution obtained at the end of each exposure is used to locate the states of the thermodynamic system representing the membrane in the energy-entropy plane (**Figure 4.3B**). Moreover, after exposure, the system is allowed to relax to equilibrium as shown by the arrows in the figure. The interaction between the system and an external field increases the energy of the system. Therefore, the higher the value of *τ_pw_* is, the higher the system’s expectation value of the energy at the end of the exposure. Interestingly, there is a much larger sudden jump in the energy value for a pulse width of *τ_pw_* = 0.5 when compared to the other pulse widths. The reason for this seen in the lipid degree of order vs. time plot predicted by SEAQT EOM (**Figure 4.3C**; arrows indicate when the PEF pulse is switched off for the respective cases). In this plot, it is noted that while the PEF minimally alters the lipid degree of order for pulse widths of *τ_pw_* = 0.125 and 0.25, the PEF pulse width *τ_pw_* = 0.5 drastically reduces the lipid degree of order and pushes the system beyond the critical point. To relate the non-equilibrium to the equilibrium, the equilibrium states obtained by the SEAQT EOM as the system state relaxes to equilibrium after PEF exposure are plotted on the figure for the lipid degree of order versus *k_B_T/J* curve obtained for the equilibrium values predicted by the Metropolis-Hastings scheme (**Figure 4.3D**). This plot shows that the membrane exposed to a single PEF pulse of *ε* = 1 for *τ_pw_* = 0.5 is brought to the same state that would be obtained by increasing the temperature past the miscibility point *k_B_T / J* > 2.3.

We next simulate the lipid membrane exposed to a single PEF for a varying scaled paramter *ϵ* (0.5, 1, and 2) and a constant exposure time *τ_pw_* = 0.25 but with the same initial condition as the case of a constant *ϵ*, i.e.,

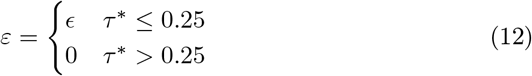

The thermodynamic state of the membrane in the energy-entropy plane for the different exposures is shown in **Figure 4.3E**. As in the case of varying values of *τ_pw_*, the higher the value of *ϵ* is, the higher the system’s expectation value of energy at the end of the exposure. There is a sudden and drastic jump in the energy value for *ϵ* = 2, which is indicative of a system that has been pushed beyond the critical point. This is validated when the lipid degree of order is traced in time (data not shown). It is seen that *ϵ* = 2 drastically reduces the order and pushes the system far beyond the critical point as shown in **Figure 4.3F**. Note that although these simulations are performed with a constant value of *J*, the observations just made are invariant for the normalized paramter *ϵ/ J*. Furthermore, two pulse trains have the same effect on the lipid membrane if the scaled parameters *ϵ/ J* and *τ_pw_* are the same. However, data from two different PEF exposures performed on two different lipid systems (e.g., from those extracted from MDA-MB-231 vs. those extracted from DCIS.com as in **Figure 1**) may not be reliably used to form a scaling law due to variance in their miscibility points. The importance of this in terms of developing a scaled PEF parameter is discussed in section 5.

#### 4.3.2. Effect of multiple pulses

To explore the impact of multiple pulses, the lipid membrane is subjected to a pulse train consisting of 3 pulses, each with *ϵ* = 1 and *τ_pw_* = 0.25. The interval between pulses is *τ*\* = 0.25. This pulse train is shown in **Figure 5A**. The lipid membrane is at the same initial state as used in the previous section and is represented by state 1 on the energy-entropy diagram (**Figure 5B**). Immediately after the first pulse, the membrane is described by state 2 which relaxes to state 3. The next pulse pushes the system across the critical point and to state 4. Following relaxation to 5, the system is pushed to a higher energy state at 6. The membrane is allowed to relax to an equilibrium distribution at 7. Note that the energy jumps due to the PEF pulse from 1 to 2, 3 to 4, and 5 to 6 are not equal. This highlights that multiple pulses induce effects on the lipid membrane that cannot be understood by a linear extrapolation of our understanding developed from a single pulse and draws attention to the unreliability of a scaling law that is based on the number of pulses. The behavior of the membrane (synthetic or living) is captured by the Ising Hamiltonian and SEAQT EOM, which show that the non-linearity/linearity of the PEF pulse number effect depends on the initial state as well as the critical miscibility point of the system.

**Figure 5:**
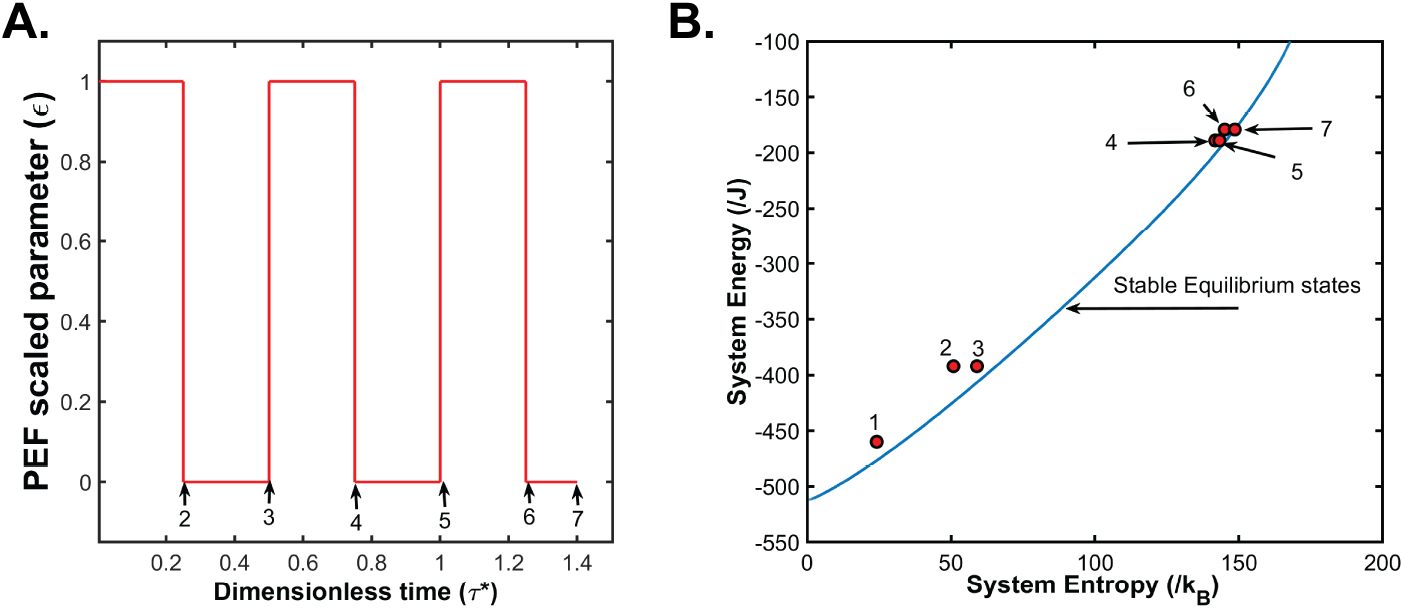
Effect of multiple PEF pulses is not additive. Multiple pulse PEF pulse train (**A**). The thermodynamic state of the lipid membrane is visualized in the energy-entropy plane (**B**). Initially at a state below the critical point (1), the lipid membrane is exposed to three pulses. The thermodynamic states corresponding to A are shown on the energy-entropy plane (**B**). Note that state 7 results from the system relaxing to equilibrium after reaching state 6.

## 5. Discussion

### 5.1. PEF induced lipid membrane poration/permeation as a phase change phenomenon

PEF induced permeabilization of the lipid membrane has been explained via an Arrhenius rate equation whereby the rate at which the pores form depends on an activation energy barrier as well as the temperature [56]. Thus, increasing the temperature increases the permeability of the lipid membrane induced by an electric field. Experiments on GPMVs/GUVs demonstrate that electric field induced pores are more likely to be formed in fluid phases rather than gel phases, i.e., the transmembrane potential required for poration in gel phases is higher than in fluid phases [57, 58]. Experiments analogous to Kraske and Mountcastle [26], which are discussed in section 2.1, are performed for electrically perturbed lipid membranes in which increasing cholesterol content in vesicle membranes reduces the electric field induced pore formation [59]. In this study, we show that cholesterol content in mammalian cells dictate their susceptibility to μsPEFs (**Figure 1**). As pointed out, such observations have also been made for nsPEFs [28]. In mammalian cells, cholesterol-rich ordered domains exist as shown in **Figure 1C**. Exposure to PEFs may induce changes in lipid order and loss of ordered domains, as have been observed experimentally using fluorescent trans-parinaric acid imaging in yeast cells exposed to a μsPEF [60]. Note that lipid peroxidation is one hypothesized mechanism behind such an observed loss of order [61]. Based on these observations, we propose to view PEF induced poration/permeation of the lipid membrane as a phase change phenomenon whereby a critical point is determined by both composition and temperature. Furthermore, observations that the lipid membrane behaves like a ferromagnet, allows us to propose the bivariate Ising Hamiltonian model in the SEAQT framework to capture the lipid dynamics perturbed by a PEF whereby the effect of the PEF is captured by a scaled parameter *ϵ*.

### 5.2. SEAQT-based experimental designs to scale the PEF parameter space

To determine the form of the scaled parameter *ϵ*, we simulate a lipid membrane represented by an Ising Hamiltonian using the SEAQT EOM. First, the eigenstructure of the Ising lattice is determined using the WLA and REN algorithms, and the SEAQT solution strategy determined via prediction of the relaxation dynamics of an isolated Ising system (**Figure 2**) is validated. We demonstrate that the SEAQT framework can reliably predict the phase changes obsereved in lipid membranes during thermal quenching (**Figure 3**). The SEAQT predictions of PEF perturbed lipid membranes are then reported. In all of these predictions, the system state is shown to evolve in a dimensionless time *τ*\* with the value of the interaction term *J* assumed constant. As discussed in section 2.2. *τ*\* is the dimensionless time defined by the ratio of the actual time and the relaxation time (*τ*\* = *t*/*τ*). In the case of lipid quenching, this relaxation time can be easily determined by comparing the experimental data to the SEAQT prediction as shown in **Figure 3**. Relaxation times for GPMVs/GUVs exposed to PEFs have been measured via fluorescence microscopy and are on the order of milliseconds to seconds [62, 8]. Comparisons between the experimental studies performed on GPMVs/GUVs and the SEAQT predictions is required for determining an appropriate choice of relaxation time and is proposed as a part of future work. The determination of the interaction term *J* for a given membrane is simple given the Ising-like behavior under thermal quenching. The miscibility temperature *T_c_* is obtainable for a given membrane. This is related to the interaction term *J* by the simple expression for criticality *k_B_T_c_/J* = 2.3.

The predictions of the PEF perturbed lipid membrane show that the phase transition can occur with variation of both *τ_pw_* and *ϵ* (**Figure 4.3**). At first glance, these two parameters would appear to be analogous to the PEF pulse width and amplitude. However, certain precautions must be taken to declare them as such. First, the value of *τ_pw_* is dependent on the relaxation time *τ* chosen. It may then be related to the pulse width (*t_pw_*) via *τ_pw_* = *t_pw_/τ*. The dimension of *ϵ* is that of energy (or the mass-length-time dimensionally of *ML*^2^*t*^−2^) whereas the dimensional form of the PEF amplitude (V/cm) is *MLt*^−3^*I*^−1^ where *I* is current. Thus, although *ϵ* is analogous to the PEF amplitude, the form of this term includes the amplitude and a term whose dimensional form is *Lt*^−1^*I*^−1^. Thus, based on this dimensional form, the additional term would involve charge and could incorporate the impedance or conductivity of the membrane with some algebraic manipulations. However, an exact form requires rigorous experiments. Furthermore, it is important to note that the value of the interaction term *J* changes with alterations in the critical temperature (GPMVs extracted from different mammalian cell lines had *T_c_* ranging from 15°C-25°C). Thus, two PEF pulses are equivalent only if they have the same value of *ϵ/ J* and *τ_pw_*. Finally, our data reveal that the effect of multiple pulses cannot be obtained by a simple addition or linear extrapolation performed on the impact of a single pulse. Furthermore, scaling terms using pulse numbers *N* (such as the scaling term ∝ *N*^0.5^ in [5]) may not be reliable since the effect of pulse is dependent on the initial state of the lipid membrane as well as the critical point. The Ising Hamiltonian and the SEAQT EOM capture the non-linearity that arises from the pulse numbers as shown in **Figure 5**.

We also speculate that the Ising model based scaling of the PEF parameter should allow the determination of the appropriate PEF-induced pore sizes in living cells. Experiments on GUVs exposed to PEFs reveal electropores that are on the order of *μm*, which is three orders higher than what is observed in living cells. All systems that follow the two-dimensional (2D) Ising universality class have the same relationship of correlation length (*ξ*) and temperature given by *ξ* = *ξ*_0_*T_c_*/(*T* – *T_c_*) where |*T* – *T_c_*| provides a distance of the system from the critical temperature. Fluorescent microscopy measurements on GUVs/GPMVs have been used in the literature [13, 12] to derive a term defined as the *structure factor*, which is akin to the magnetic susceptibility *χ* in the Ising model. The *structure factor* behaves as given by *χ* = *χ*_0_*T_c_*/(*T* – *T_c_*)^7/4^. Using the concept of scaling the existence of nanometer sized ordered domains in living cells have been explained from observations made on GUVs where only *μ*m-sized domains are seen. For example, a measured correlation length of 1 *μ*m at a temperature of 23.7°C that is a slightly above (0.3°C) the critical temperature of 23.4°C for a GUV can be extrapolated to provide an estimate for a living cell membrane if one accounts for the difference in the physiological temperature (37° C) and critical temperature 23.4°C. A direct scaling can be performed to estimate the correlation length to be *χ* = 1*μ*m ×0.3°C / 13.6°C = 22 nm at the physiological temperature. Such a scaling approach can be taken to provide approximations of pore sizes induced by different PEF pulse trains.

Finally, once a form of the scaled PEF parameter *ϵ* is known, experiments with only one type of PEF pulse train (e.g., μsPEF or ms-PEF) will be enough to derive information about another PEF parameter space. Furthermore, the equivalent pulses should always induce comparable effects on the plasma membrane regardless of whether they are on a synthetic or living cell membrane. This can be powerful, since the mechanistic understanding developed in one space (e.g., peroxidation effects using nsPEF) can be extended to another (e.g., H-FIRE).

### 5.3. Extension of the scaling law to other cellular compartments

The inclusion of transmembrane protein and actin cytoskeleton interaction influences the rate of electric field induced pore formation on the lipid membrane [63, 59]. For example, experiments on PEF exposed GUVs with actin networks have smaller pores and larger resealing times when compared to those GUVs without actin networks [62]. Interactions between the cytoskeleton and the lipid membrane are important in restricting phase separation and have been modeled as an Ising model with the cytoskeleton created as Voronoi structures on the Ising lattice [64]. While such a mathematical representation has not been modelled here, the SEAQT framework could be used to do so as could the kinetic Monte Carlo strategy, which we recently implemented for PEF induced integrin clustering [65]. Interactions of the actin cytoskeleton with the the lipid membraneare not only important for PEF-induced pore formation but also for downstream effects on the organelles [66, 67]. Thus, a scaling strategy with actin cytoskeleton interactions may be possible in the future.

Computational challenges for such approach with the SEAQT is of course to conceive a statistical approach to obtain the eigenstructure. The non-Markovian techniques such as those introduced here may be deployed for such information. Furthermore, another challenge in solving systems with substantial differences in magnitude in both the degeneracies and the probabilities is that the system becomes highly stiff. Due to the non-sparse nature of the Jacobian in solving the SEAQT EOM, we eliminated Jacobian-free Newton-Krylov as an option. We believe the extension to highly stiff larger systems may involve using multi-grid methods to solve for a non-sparse Jacobian or alternatively may require using the Hilbert subspace approach proposed by Li and von Spakovsky [23, 24], and is left for future work.

## 6. Conclusion

PEFs offer an attractive modality for cancer therapeutics and tissue regeneration. However, the boon of achieving different cellular responses by altering PEF parameters is also the bane for this modality due to the vast parameter space. This space represents a bottleneck to exploring mechanistic questions as to how PEFs influence cell behavior. We provide a solution to this conundrum via the development of scaling laws based on thermodynamics and a system Hamiltonian. We demonstrate the approach for lipid membranes and provide insights for future experiments and computational efforts.

## 7. Acknowledgements

We would like to thank Professor Stephen Eubank (University of Virginia, U.S.A.), Dr. Yihui Ren (Brookhaven National Laboratory, U.S.A.), and Dr. Madhurima Nath for their discussions on the WLA and REN algorithms.

## 8 Author contribution

Project plan and study design: I.G., S.S.V. and M.R.v.S.

Coding and simulations: I.C.

Experiments: I.C. and R.B.

Data analysis and manuscript writing: I.C., R.B., S.S.V., and M.R.v.S.

## Appendix A. Appendix name

### Appendix A.1. PEF setup

**Figure A.1:**
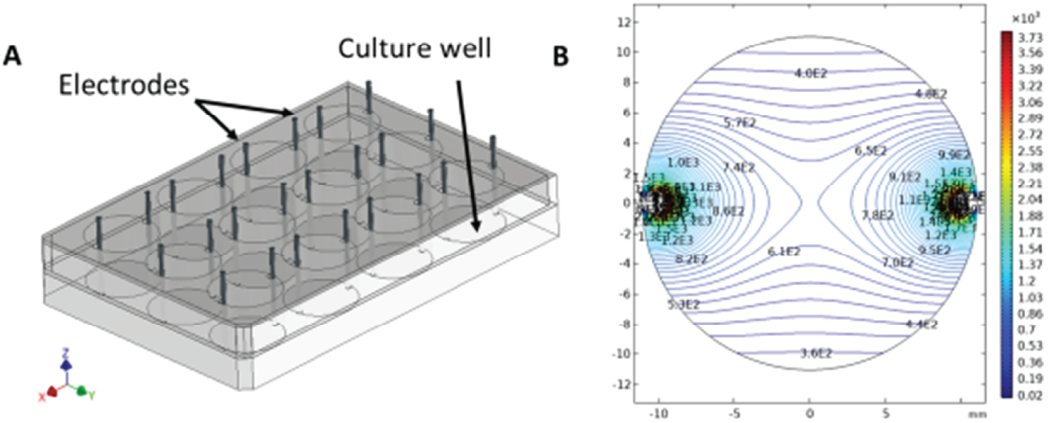
Illustration of the experimental setup for delivering the μsPEF **(A)**. Prediction made via a finite element simulation in the COMSOL software. The distribution of the electric field, as predicted by COMSOL, is shown in **(B)**. Contour lines connect areas of the same electric field strengths.

### Appendix A.2. WLA and REN algorithm details

#### Appendix A.2.1. Wang-Landau algorithm (WLA)

The WLA is a histogram technique, proposed by Fugao Wang and David P. Landau [52, 53], to extract information about the degeneracy associated with energy eigenlevels. As opposed to the unbiased random walker approach, which is more likely to visit energy eigenlevels that have higher degeneracies, the WLA adopts a Metropolis-Hastings based sampling whereby a random walk is accepted if the move is towards an energy level that has been sampled less times than the current one. Mathematically, a transitional probability *p_i→j_* (from state *i* to *j*) is defined in the same manner as that for the Metropolis-Hastings algorithm such that 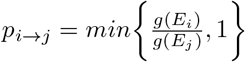 where the term *g*(*E_k_*)(*k* = *i,j*) is the total number of times the random walker has accessed the *k^th^* energy eigenlevel. If the transitional probability is greater than a random number drawn from a uniform distribution, the random walker’s move is accepted. Every time a move is accepted, the term *g*(*E_accepted_*) is multiplied by a modification factor *f*, and a histogram count *h*(*E_accepted_*) is increased by one. The WLA starts with a value *f* = *e*^1^ for the modification factor and reduces it by half every time a histogram flatness (e.g., 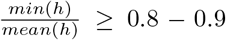) is reached. The histogram is reset when the value of the modification factor changes, while the total count *g* is retained. The algorithm is run until the value of *f* ≈ 0 (practically to a tolerance value such as 10^-8^). Note that the WLA is non-Markovian although it adopts a transitional probability that has a form similar to that of the Markovian Metropolis-Hastings algorithm. The WLA biases the random-walker according to the recorded number of times eigenlevels have been accessed, and, therefore, the history of the random walker dictates the convergence rate.

#### Appendix A.2.2. Ren-Eubank-Nath (REN) algorithm

Using a WLA to obtain energy eigenlevel and degeneracy information for the bivariate case is very challenging since it entails performing random walks in two dimensions, one for the energy and the other for the lipid degree of order. In other words, a Monte Carlo scheme to track both the energy and configuration of the system must be devised. Alternately, the approach proposed by Ren, Eubank and Nath (REN) can be used. It is based on graph theory and constructs a parallel scheme to estimate bivariate distributions in a fashion inspired by the Moore-Shannon network reliability approach. Briefly, the work by Ren *et al*. [54] expresses the partition function of the bivariate Ising Hamiltonian in the same form as the reliability of an interaction network consisting of vertices and edges such that

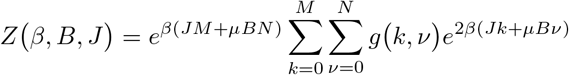

where *g*(*k, ν*) is the degeneracy associated with a state with *k* discordant edges and *ν* the the number of spins up. *J* and *B* have the same meaning as those used in **Eq. 1**, and *β* = 1/*k_B_T*. Moreover, the bivariate degeneracy *g*(*k, ν*) is related to the conditional probability *p*(*k|ν*) of finding a given discordant edge number *k* if the number of spins up is *ν*: 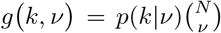. In this relationship 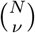 is the combination of *N* nodes taken *ν* at a time. Thus, Ren *et al*. [54] propose estimating the conditional probability *p*(*k|ν*) using a Monte Carlo technique that incorporates Kawasaki spin-exchange (i.e., swapping spin locations rather than spin flips to conserve the ∑_*i*_ *σ_i_* to a given number). By its inherent design, the REN algorithm unlike the WLA is parallel since one may run simulations for different values of ∑_*i*_ *σ_i_* simultaneously. However, the REN does encounter the same issues as in a naïve Monte Carlo technique to correctly predict the probability of energy eigenlevels with lower degeneracies especially towards the tail of the energy spectrum. To counter this, Ren *et al*. suggest adopting a similar strategy to WLA. In this approach, a transitional probability (*p_i→j_*) from a state defined by the discordant edge number *i* to another state *j* but having the same number of spins up (or ∑_*i*_ *σ_i_* = constant) is defined as 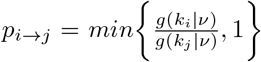. The choice of this transitional probability allows the random walker to access states that would otherwise be very difficult to access. Just as with the WLA, a modification factor, *f*, whose value is chosen to be *e*^1^ at the start of the simulation, is reduced by half every time the least value in the histogram is 1 after a certain number of Monte Carlo sweeps. The histogram is reset when the value of the modification factor changes, while the total count g is retained. The algorithm is run until the value of *f* ≈ 0 (e.g., to a tolerance value such as 10^-6^). However, the REN algorithm uses the histogram in a different way than the WLA since just as with the WLA, there can be a debate on how to decide the appropriate number of Monte Carlo sweeps upon which the histogram criterion is checked.

During the implementation of REN for this work, it was observed that although one needs to only run a simulation for spin numbers *ν* = 3 through *N*/2, the geometries associated with the lowest discordant edge numbers for *ν* = 3 through *N*/4 have a distinct spatial pattern compared to those associated with the lowest discordant edge numbers for *ν* = *N*/4 + 1 through *N*/2. Starting the random walk in geometries associated with the former lowest discordant edge numbers drastically improves the convergence rates of the algorithm (highlighting the non-Markovian nature of the algorithm). Similarly, for *ν* = *N*/4 + 1 through *N*/2, starting the random walk alternatively at geometries associated with the highest and lowest discordant edge numbers allows faster convergence.

### Appendix A.3. Initial state preparation and SEAQT simulation details

A non-equilibrium state of the system is prepared by generating a probability profile *F*(*ϵ*) where the variable *ϵ* is the energy eigenlevel. A profile could be prepared by a single Gaussian function or the superposition of multiple Gaussian functions. A vector 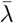 is then determined such that the following conditions hold:

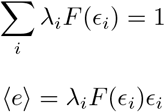

These conditions make sure that the probabilities associated with each energy eigenlevel sum to unity and that the value of the expected system energy 〈*e*〉 is chosen such that the probability distribution associated with that value at stable equilibrium is known a priori from the WLA prediction. Upon solving for the vector 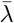, the probability for each eigenlevel may be obtained, i.e., *p_i_* = λ_*i*_ *F*(*ϵ_i_*). An example of a non-equilibrium state prepared using the aforementioned approach is shown in **Figure 2C** in which a univariate distribution at dimensionless time *t*\* = 0 is shown for a 50×50 lattice.

To prepare a bivariate distribution, i.e., construct a distribution of both energy and lipid degree of order, the univariate distribution obtained from the aforementioned methodology is multiplied with another function *G*(*k, ϕ_i_*), which is a function of the discordant edge number *k* and the lipid degree of order *ϕ_i_* such that the summation of probabilities along a discordant edge is unity, i.e.,

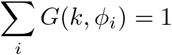

The function *G*(*k, ϕ_i_*) may assume the shape of a Gaussian along a discordant edge number *k*. An example of a non-equilibrium state prepared using this approach is shown in **Figure A.2A** in which a bivariate distribution at dimen-sionless time *t*\* = 0 is shown for a 16×16 lattice. Another set of non-equilibrium states prepared using this approach for PEF perturbed lipid membrane is shown in **Figure A.3**.

**Figure A.2:**
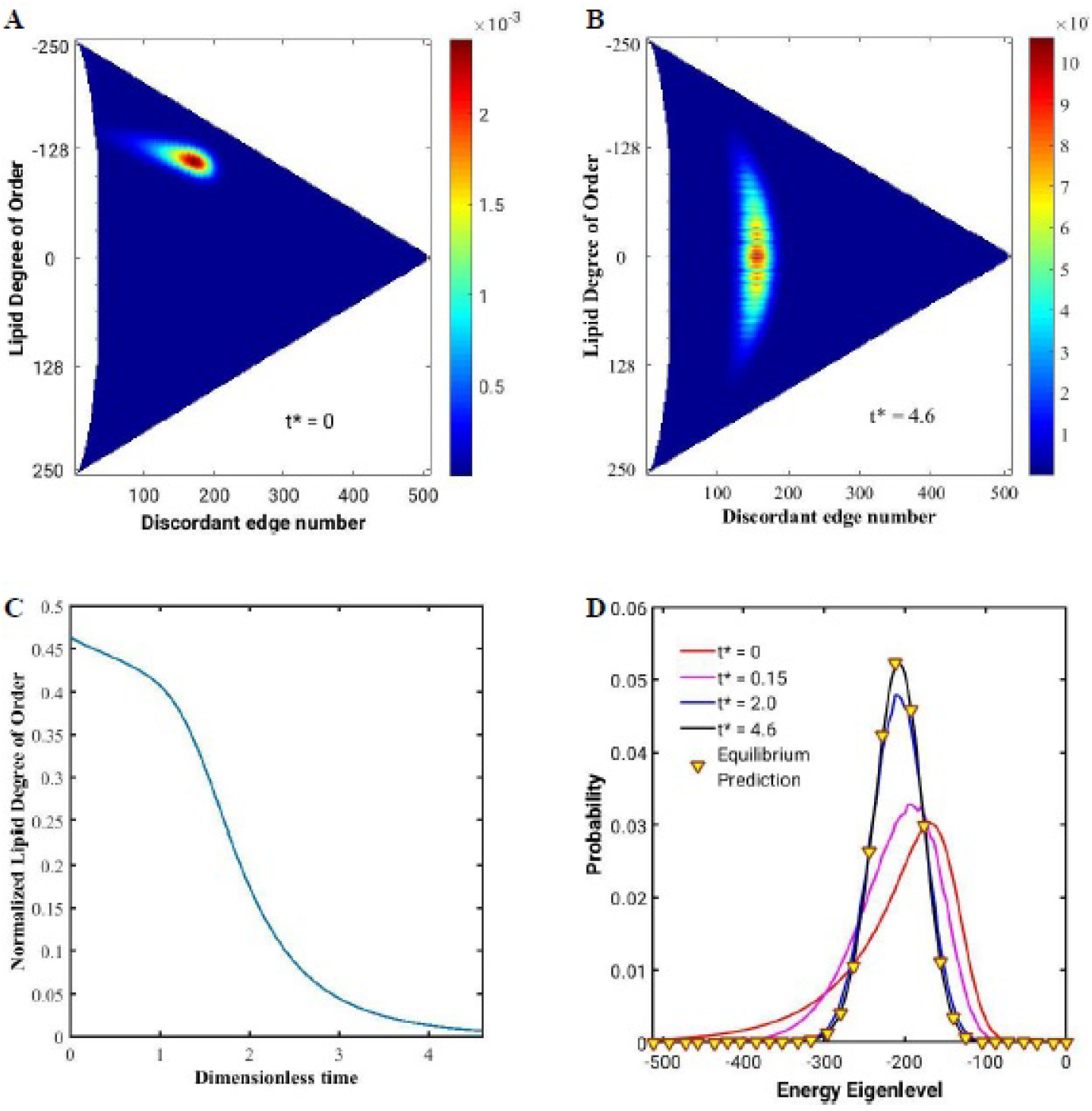
SEAQT prediction of the temporal evolution of the bivariate distribution. The bivariate distribution of probabilities prepared as the initial condition (*t*\* = 0) is shown for a 16×16 lattice system in (**A**). Note that the preparation is made such that the expected system energy is associated with a stable equilibrium temperature that is higher than the non-dimensional critical temperature (kT/J ~2.3). The system relaxes to a bivariate distribution (*t*\* = 4.6) that is associated with an expected lipid degree of order of 0 (**B**). The temporal variation of the absolute value of the normalized (i.e., divided by *N* = 256) lipid degree of order is shown in (**C**). The temporal evolution of the univariate distributions obtained by summing over the discordant edge numbers in the bivariate distribution is shown in (**D**).

The system of equations represented by **Eq. 9** is solved using the MATLAB suite of ordinary differential equation (ODE) solvers. For a 16×16 lattice, a Runge-Kutta integration scheme implementation, ODE45, is used. Note that the system represented by **Eq. 9** for larger systems, such as the 50×50 lattice, becomes highly stiff due to the order differences in the values of degeneracies and probabilities. Thus, the stiff-solver ODE15s, based on an implicit scheme, is used. Moreover, for the bivariate cases in this chapter, a lattice of 16×16 is used. This is because with increasing lattice size the differences in magnitude in both the degeneracies and the probabilities in the SEAQT EOM also increase, thus, making the system highly stiff. Even a moderately sized lattice such as a 32×32 grid requires over 1 TB of RAM in order to be solved using the ODE15s solver. The extension of the bivariate predictions to larger lattices is left for future work and may involve using multi-grid methods to solve for a non-sparse Jacobian or alternatively may require using the Hilbert subspace approach proposed by Li and von Spakovsky [23, 24].

As to convergence to stable equilibrium, this is detected via three checkpoints. First, the derivatives of the probabilities with respect to time are calculated. If the time rates of change of all system probabilities are negligible (e.g., ≤ 10^-4^) the system is acceptably close to stable equilibrium. Moreover, the variation of entropy with respect to time can also be monitored (the rate of change of entropy Δ*s* ≥ 0). The third checkpoint is to compare the predicted probability distribution to that predicted by the WLA for the given value of expected system energy.

**Figure A.3:**
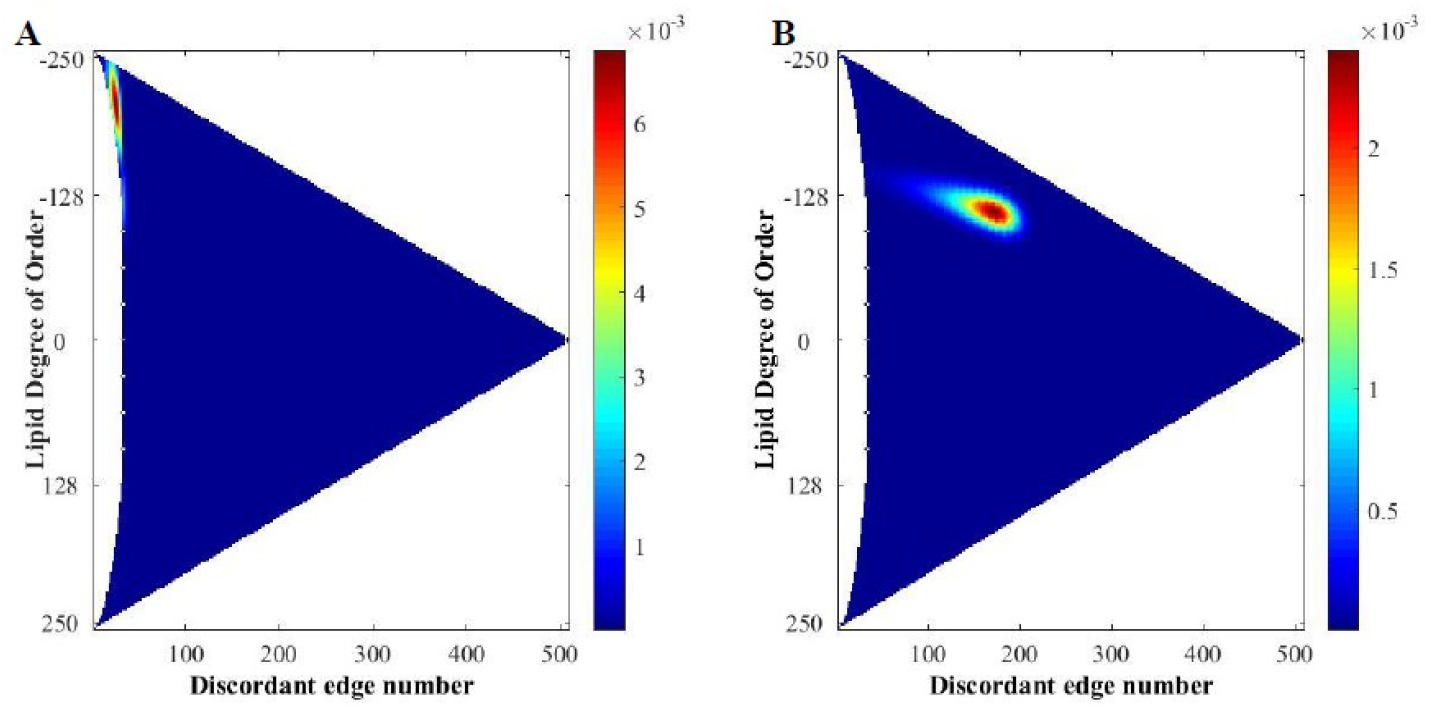
Two representative initial conditions tested here. The membrane can be below the critical point where the non-zero lipid order is favored (**A**) or above the critical point where it is not favored (**B**).

## Footnotes

1 Note that the fate of pores depends on the energy delivered via the PEF, and hence any of the PEF parameters (i.e., pulse number, width, and amplitude) may dictate pore resealing dynamics. For simplification, in this paragraph we discuss only when amplitude is altered in the μsPEF pulse train.

